# Deep Learning Enables Automated Segmentation and Quantification of Ultrastructure from Transmission Electron Microscopy Images

**DOI:** 10.1101/2025.11.05.686793

**Authors:** Anqi Zou, Winston Tan, Jiayi Ji, Florencia Rojas-Miguez, Laura Dodd, Emily Oei, Sarah R. Vargas, Stephen P. Berasi, Hongying Yang, Hui Chen, Joel M. Henderson, Xueping Fan, Weining Lu, Chao Zhang

**Author notes:** Corresponding authors E-mail addresses (C. Zhang), (W. Lu), (X. Fan). These authors contributed equally to this work (Anqi Zou, Winston Tan, Jiayi Ji).

## Abstract

Transmission electron microscopy (TEM) has become an essential technique for observing subcellular ultrastructure, and is widely used in both clinical diagnosis and biomedical research. However, analysis of TEM data remains extremely labor-intensive and often inconsistent across operators due to the lack of dedicated computational methods. Here, we present TEAMKidney, a deep learning framework for accurate and scalable measurement of ultrastructures in TEM images across species, magnifications, and instrument platforms. We collected 12,991 TEM images from patients with multiple kidney diseases and from different animal models. By combining a self-training-based semantic segmentation stage with a TEM-tailored panoptic segmentation model, we address two major challenges in TEM data analysis: the lack of accurately labeled training data and the difficulty of achieving high segmentation accuracy for complex ultrastructure. Application of TEAMKidney to both human and animal images successfully reveals disease-associated changes in two critical glomerular ultrastructures: the glomerular basement membrane and podocyte foot processes. In addition to significantly outperforming existing tools, TEAMKidney shows close agreement with pathological expert measurements used in clinical assessment protocols. By reducing dependence on manual tracing while preserving expert-level accuracy, TEAMKidney demonstrates that deep learning can substantially reduce the burden of image analysis in both clinical pathology and biomedical research settings.

## Introduction

Transmission electron microscopy (TEM) provides high-resolution visualization of cellular ultrastructure beyond the resolving capabilities of light microscopy, immunofluorescence (IF) staining, immunohistochemistry, or molecular assays^1, 2^ (Fig 1A). Beyond its extensive use in biomedical research and drug discovery, TEM remains an important diagnostic tool in many clinical areas, including infectious diseases, oncology, ophthalmology, neurology, and especially renal pathology^3-5^.

**Fig. 1.**
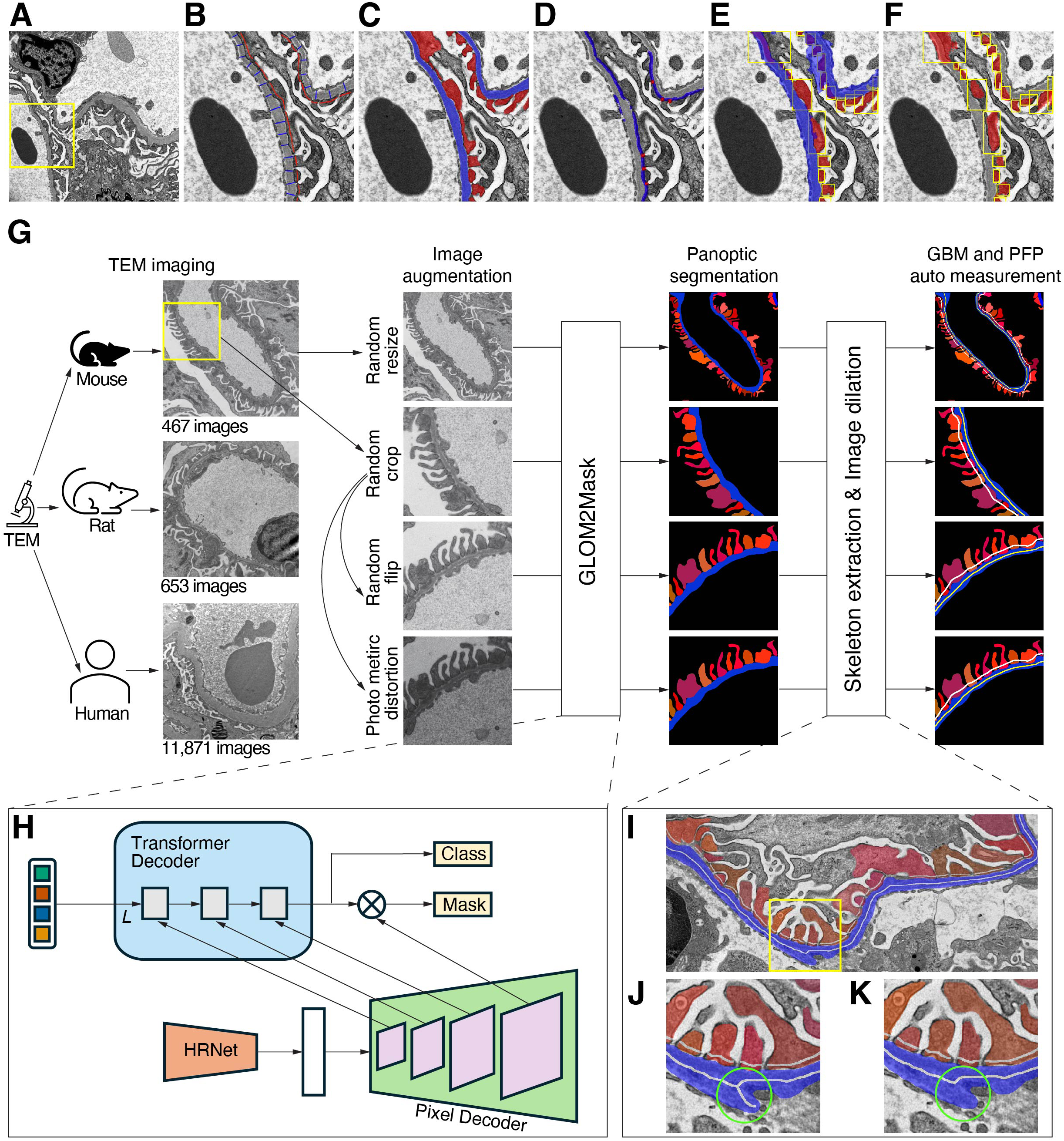
Lack of existing models for accurate TEM image analysis and overview of the proposed framework. (A) Example TEM image of a podocyte-specific ILK conditional knockout mouse. (B) The boxed region of the glomerular filtration barrier in (A) with manually measured PFP in red and GBM in blue. (C) Ground truth segmentation of PFP and GBM. Segmentation results obtained using (D) FPW-DL, (E) SAM, and (F) MedSAM models. (G) Workflow of the proposed TEAMKidney framework for augmentation, segmentation, and quantitative analysis of mouse, rat, and human TEM images. (H) The proposed Glom2Mask network architecture for panoptic segmentation. (I) Example image showing post-segmentation of GBM and PFP. (J) Boxed region in (I) illustrating the initial skeleton extraction. (K) Demonstration of skeleton extraction error correction by the post-segmentation analysis algorithm.

Chronic kidney disease (CKD) affects over 37 million people in the US^6^ and 850 million worldwide^7, 8^. It has been recognized as a leading public health burden in the US. By 2027, the direct healthcare costs for kidney diseases in the US were projected to reach $173.4 billion^9^. Diabetic kidney disease and glomerular disease account for more than 50% of CKD cases^10, 11^ and are the leading cause of kidney failure^12, 13^. These disorders encompass a broad spectrum of conditions characterized by distinct patterns of glomerular ultrastructural damage, which are associated with different therapeutic strategies and clinical outcomes. In the past decade, TEM has become an essential and often decisive component of the diagnostic modalities for kidney diseases^14^. Some major nephrology and pathology centers now utilize TEM as a routine diagnostic tool, applying it to more than 80% renal biopsies^14, 15^.

Despite its value, significant barriers limit the broader adoption of TEM. The technique remains highly labor-intensive^14^ and producing a full pathology report can take up to 26 working days on average^5^, primarily due to the need for expert renal pathologists to analyze the ultrastructure, such as glomerular basement membrane (GBM) and podocyte foot processes (PFP). Measurement of GBM and PFP widths is widely used for both research and clinical diagnosis, yet researchers and renal pathologists still rely primarily on manual measurement of TEM images (Fig 1B). This process is time-consuming and restricts the advancement of CKD research and clinical diagnosis.^16-19^. Manual measurement also introduces significant inter-operator variation and bias^20-22^. Due to the inherent characteristics of TEM data, such as grayscale-only imaging, low contrast, and fuzzy boundaries between ultrastructural features, which make conventional artificial intelligence (AI) models struggle to delineate fine details accurately. Currently, no available tools achieve results as accurate as manual labeling (Fig 1C).

During the past decade, semi-automated methods have been piloted for GBM and PFP measurements^20, 23, 24^ (Table 1). However, these methods have suffered from low accuracy and still require expert user intervention for each image. In parallel, several fully automated deep learning-based approaches have been proposed for TEM image analysis, targeting the segmentation of GBM^25, 26^, PFP^21^, or both^27-29^, as well as other related ultrastructural components^25, 30, 31^. Most of these methods adopt U-Net-based semantic segmentation frameworks. While such models demonstrate improved efficiency over traditional pipelines, they are inherently limited in their ability to distinguish individual PFP instances, a critical requirement for accurate morphometric analysis. Moreover, many of these studies are trained and evaluated on single-center or disease-specific datasets, limiting generalizability across species, disease types, and imaging protocols^25, 26, 28^. In addition, suboptimal segmentation accuracy further constrains their applicability for precise downstream ultrastructural measurements. More recently, the automated model FPW-DL was developed to measure PFP width^31^, with publicly released code and a valuable human disease TEM dataset. The method demonstrates stable performance when applied to human samples during testing. However, its reliance on identifying slit diaphragms as proxies for PFP width measurement imposes intrinsic methodological limitations, leaving substantial room for improvement in measurement accuracy (Fig. 1D). The Segment Anything Model (SAM)^32^, the recently published model, has become one of the most successful deep learning models for general image segmentation. A variant fine-tuned for medical images, called MedSAM^33^, has also been released. While both models outperform most others in many biomedical image segmentation tasks, their performance in TEM image analysis is unreliable, even when assisted with manual prompting (Yellow boxes in Fig 1E, 1F). The need for precise prompts underscores the challenge of applying SAM to TEM images. Building upon these developments, a recent study proposed TEM-AID^34^, a multi-stage TEM analysis framework integrating detection, prompt-based SAM, measurement, and classification. However, its reliance on sequential processing and human-in-the-loop refinement increases system complexity and limits scalability, and its quantitative analysis primarily focuses on GBM, with limited support for simultaneous PFP quantification. Significant variance across different diseases, the lack of high-quality labeled training data, high resolution images with multi-scale and fine-grained features, and boundary ambiguity make TEM image analysis extremely challenging for general AI models. Together, these challenges highlight the urgent need for a specialized digital pathology platform capable of efficient and precise TEM image analysis.

**Table 1.** Data Collection and Selection Criteria.

| Group | Total | Manually labeled | Manually measured | Training & Validation | Evaluation |
| --- | --- | --- | --- | --- | --- |
| Mouse models |  |  |  |  |  |
| WT | 107 | 48 | 20 | 48 | 61 |
| ILK cKO | 241 | 67 | 20 | 67 | 174 |
| Col4A3 | 93 | 0 | 20 | 0 | 93 |
| Rat models |  |  |  |  |  |
| WT | 69 | 0 | 20 | 0 | 69 |
| PHN | 584 | 540 | 20 | 540 | 44 |
| Human models |  |  |  |  |  |
| Normal | 1361 | 71 | 20 | 0 | 1361 |
| Fabry males | 4301 | 76 | 20 | 468 | 3833 |
| Fabry females | 2487 | 40 | 20 | 0 | 2487 |
| DKD | 2463 | 33 | 20 | 0 | 2463 |

In response to these challenges, we introduce TEAMKidney, a novel deep-learning framework designed for precise segmentation and quantification of TEM ultrastructure. Through extensive generation, collection, and curation of approximately 13,000 TEM images representing various kidney diseases across human, mouse, and rat samples, TEAMKidney is the first deep learning model capable of robustly analyzing TEM images across different magnifications, disease states, and species. To address the challenge of limited high-quality labeled data, in addition to having experienced pathologists meticulously annotating the ultrastructure of glomeruli in 315 TEM images, we incorporated a self-training strategy to leverage large amounts of unlabeled data. Furthermore, our specifically designed panoptic segmentation model can accurately distinguish individual ultrastructural components even in the absence of clear boundaries, which represents a major challenge for existing methods.

To validate the framework, we conducted extensive quantitative evaluations. In addition to comparisons with expert manual annotations, TEAMKidney significantly outperforms existing methods. Robust performance across data from different sources and species demonstrates the strong generalizability of TEAMKidney. Successful application to both animal models and patient datasets with diverse disease states further indicates its ability to capture subtle ultrastructural changes in real-world TEM data. Overall, this approach has the potential to enhance diagnostic accuracy, facilitate large-scale research on kidney disease, and support clinical decision-making in nephrology. Moreover, it can readily be extended to other clinical domains, making it a promising step toward practical deep learning-based analysis for general TEM imaging.

## Results

### The Overview of the TEAMKidney Framework

TEAMKidney is a deep learning-based automated framework for TEM image analysis, designed to precisely quantify ultrastructure, such as GBM and PFP widths (Fig. 1G). The framework comprises three key components: (1) integrating HRNet with self-training for initial semantic segmentation (Fig. 1H), (2) developing a novel model for panoptic segmentation of glomerular ultrastructure (Fig. 1H), and (3) performing post-segmentation analysis to robustly quantify detailed ultrastructure, such as GBM width and PFP width (Fig. 1I-K).

To train and validate our framework across diverse pathological conditions and species, our study conducted an analysis of both animal and human models under normal and diseased states. For the mouse, the *Integrin linked kinase* podocyte-specific conditional knockout (ILK cKO)^35, 36^ was used for model training, whereas the type IV collagen subunit alpha 3 knockout (Col4a3 KO) model was reserved as an independent evaluation dataset to assess model generalizability. For rats, the passive Heymann nephritis model (PHN)^37^ was used, which was originally developed to characterize Membranous nephropathy (MN). For human samples, we focused on the comparison among healthy donors, diabetic kidney disease (DKD)^38, 39^ and Fabry disease^40, 41^. For each species and disease model, a corresponding normal control dataset was included to enable both training and validation.

This framework was implemented to automatically segment, label, and measure ultrastructure in TEM images, thereby eliminating the need for manual tracing and reducing observer variability. This design allows the framework to address diverse measurement tasks, including interspecies comparisons, disease-associated ultrastructural changes, and quantitative evaluation of experimental and clinical datasets.

### Two-Stage Segmentation Model for Improved Fine Ultrastructure Recognition

Due to the high magnification and resolution of grayscale TEM images, ultrastructural features often appear irregular and have unclear boundaries. In addition, TEM images are acquired across multiple magnification levels and spatial resolutions, resulting in substantial scale variation of ultrastructural components within and across datasets. These challenges prevent the direct adaptation of existing segmentation software for TEM image analysis (Fig. 1D-F)^31-33^. To address this, we design a novel deep learning model, Glom2Mask, that fuses both semantic and panoptic segmentation approaches, enabling precise segmentation of ultrastructural features across species and imaging scales.

In the semantic segmentation stage (Fig. 2A), we evaluated three backbone networks for comparison: DeepLabV3^42-44^, PSPNet^45^, and HRNet^46^. All three models are effective in capturing global context and fusing multi-scale features. Overall, the models achieved comparable performance (Fig. 2B). Among them, HRNet attained the highest Dice scores across most categories and demonstrated markedly greater training efficiency. Although the average Dice scores of the three methods were similar, HRNet exhibited the most robust and consistent performance. Therefore, we selected HRNet as the backbone for our panoptic segmentation model in the second stage (Fig. 2C). In contrast, DeepLabV3 often failed on low-quality images (Fig. 2C, Supplementary Fig. 1), resulting in unreliable segmentation, while PSPNet tended to misclassify unrelated ultrastructure (Fig. 2C, Supplementary Fig. 1), particularly mistaking epithelial cells on glomerular capillaries for foot processes.

**Fig. 2.**
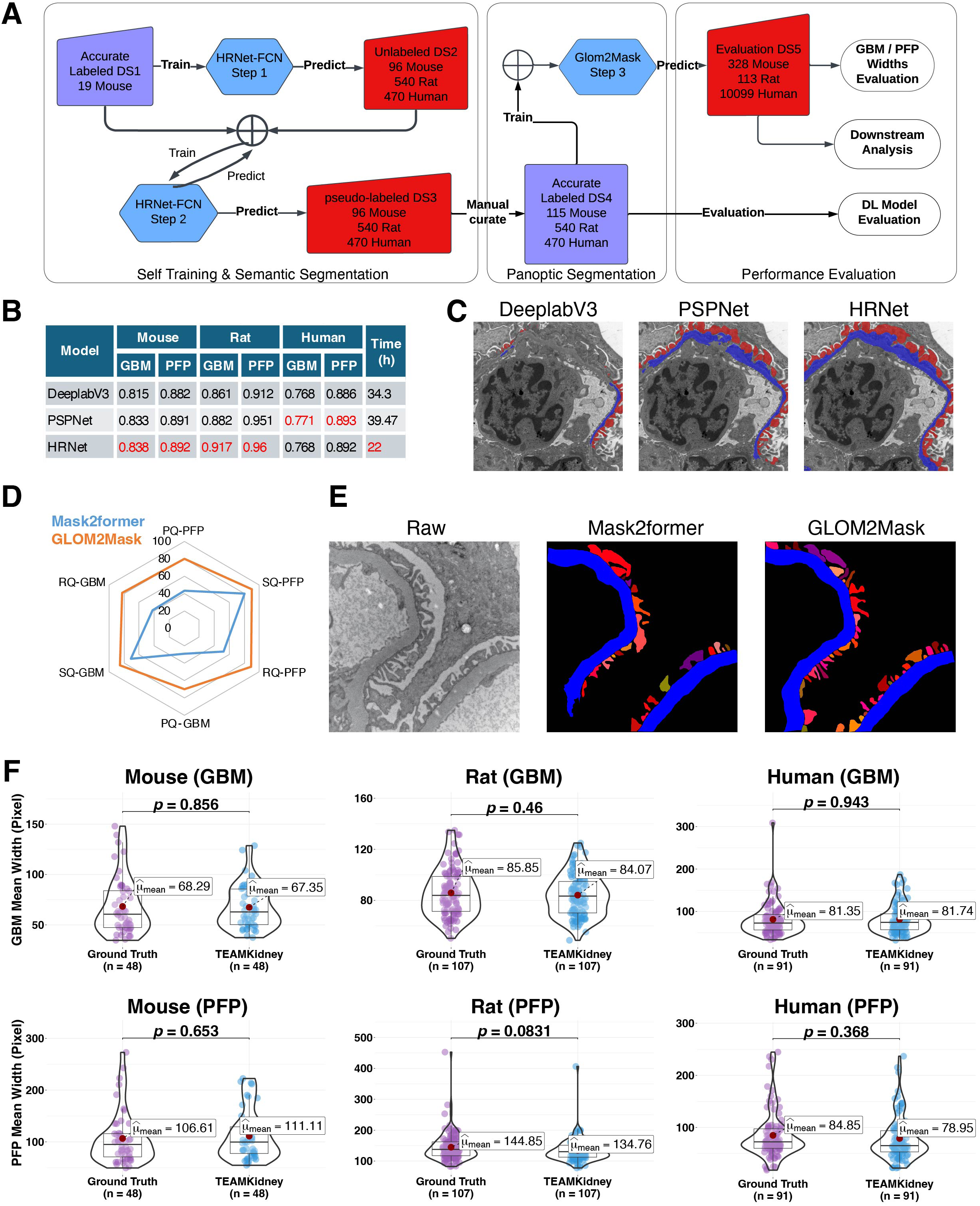
Detailed structure and evaluation of the TEAMKidney framework. (A) Workflow of the segmentation stage incorporating a self-training strategy. (B) Performance and training time comparison of different backbone networks for semantic segmentation across datasets. (C) Representative TEM images from Rat samples segmented by the three backbone networks for qualitative comparison. (D) Performance and training time comparison between the proposed Glom2Mask model and the benchmark Mask2Former for panoptic segmentation. (E) Representative image showing segmentation results from Glom2Mask and Mask2Former. The GBM is segmented as a stuff class (blue), and PFP as thing instances (various shades of red). (F) Comparison of GBM and PFP measurements between the TEAMKidney algorithm and ground truth across different species.

Although our semantic segmentation model accurately recognized major ultrastructure, direct post-segmentation analysis to identify individual instances, such as PFP and slit diaphragms, remained highly challenging and often inaccurate, similar to most existing models^28, 29^. To overcome this limitation, we introduced a panoptic segmentation stage (Fig. 1H) described in our proposed Glom2Mask model. This architecture utilizes the extracted features from HRNet trained in the first step. In addition, using annotations generated during the semantic segmentation stage, along with manually refined corrections, we constructed an expanded and higher-quality labeled training dataset to improve panoptic segmentation accuracy.

The panoptic segmentation performance of the proposed Glom2Mask and the baseline Mask2Former models were compared using standard panoptic segmentation metrics: Panoptic Quality (PQ), Segmentation Quality (SQ), and Recognition Quality (RQ), evaluated for both things (PFPs) and stuff (GBM) classes. The results show that Glom2Mask outperformed the baseline Mask2Former (Fig. 2D), achieving higher PQ, SQ, and RQ scores for both PFP and GBM while requiring fewer training iterations (66,000 vs. 360,000). The quantitative results corresponding to Fig. 2D are summarized in Table S1. These findings demonstrate superior segmentation accuracy and training efficiency of the proposed model. Fig. 2E shows representative segmentation results from Glom2Mask and Mask2Former. The visualizations demonstrate that Glom2Mask accurately distinguishes continuous GBM structures (blue) from discrete PFP instances (various shades of red), further supporting its robustness and generalizability across species.

To evaluate the robustness of our method to variations in image resolution, which may mimic the differences in images generated by different institutes or instruments, we conducted a controlled downsampling experiment using TEM images at multiple scales. As shown in Supplementary Fig. 2A-B, representative segmentation results and corresponding input images demonstrate that meaningful GBM and PFP boundaries can already be identified at moderate downsampling levels, with progressively improved delineation as resolution increases.

Quantitative analysis further confirms this observation (Supplementary Fig. 2C), showing that Dice scores for both GBM and PFP increase rapidly with higher resolution and reach a stable plateau beyond a downsampling ratio of approximately 0.1. Notably, segmentation performance remains consistently high across a broad range of resolution scales, indicating that the proposed method is robust to substantial resolution degradation and generalizes well across varying imaging conditions.

### Concordance of TEAMKidney Predictions with Ground Truth Across Species

To evaluate how segmentation accuracy affects ultrastructure measurement, we generated ground truth labels for a validation dataset comprising 48 images from 8 mouse glomeruli, 107 images from 12 rat glomeruli, and 91 images from 5 human glomeruli. All validation images were excluded from model training. GBM and PFP annotations were manually labeled by three human experts to generate precise ultrastructure masks. Then, the post-segmentation quantification algorithm was applied to both the TEAMKidney predictions and the expert-generated ground truth masks in this validation dataset. Quantified GBM and PFP width values obtained from both ground truth labels and TEAMKidney predictions were compared across mouse, rat, and human samples at the individual image level. No statistically significant differences were observed between the ground truth and TEAMKidney measurements for either GBM or PFP width across all three species (Fig. 2F), demonstrating strong concordance and reliability of the proposed framework. These results indicate that our method provides highly reliable GBM and PFP width measurements across diverse datasets.

### Distinguishing Ultrastructural Changes in Multiple Animal Models of Kidney Disease

Animals, especially rodents, are widely used to model a variety of kidney diseases. Ultrastructural changes reflect disease progression and serve as indicators of treatment efficacy during drug development^47^. To test the robustness of our framework for detecting pathophysiological changes in the glomerular filtration barrier, we applied our automated method to measure two well-established mouse models and a rat model of glomerular diseases (Fig. 3A, B). In the mouse models, we used Col4a3 KO, an animal model of Alport syndrome^48^ and ILK cKO mice, an animal model of focal segmental glomerulosclerosis (FSGS), to represent two distinct mechanisms of kidney glomerular injury^35, 36, 48^. Col4a3 KO mice exhibit severe GBM disorganization due to the lack of type IV α3 collagen, which is a crucial component of the mature GBM, whereas ILK cKO mice develop podocyte detachment and cytoskeletal abnormalities with the absence of ILK protein, which is required for podocyte adhesion to the GBM (Fig. 3A, C). Consistent with these pathological features, our automated analysis revealed significantly increased GBM widths in Col4a3 KO mice compared with wild-type (WT) controls. Similarly, PFP widths were markedly widened for the ILK cKO mouse model. These quantitative differences align with the expected ultrastructural alterations associated with abnormal GBM of the Alport mouse model and podocyte injury in the ILK cKO FSGS mouse model (Fig. 3C, D).

**Fig. 3.**
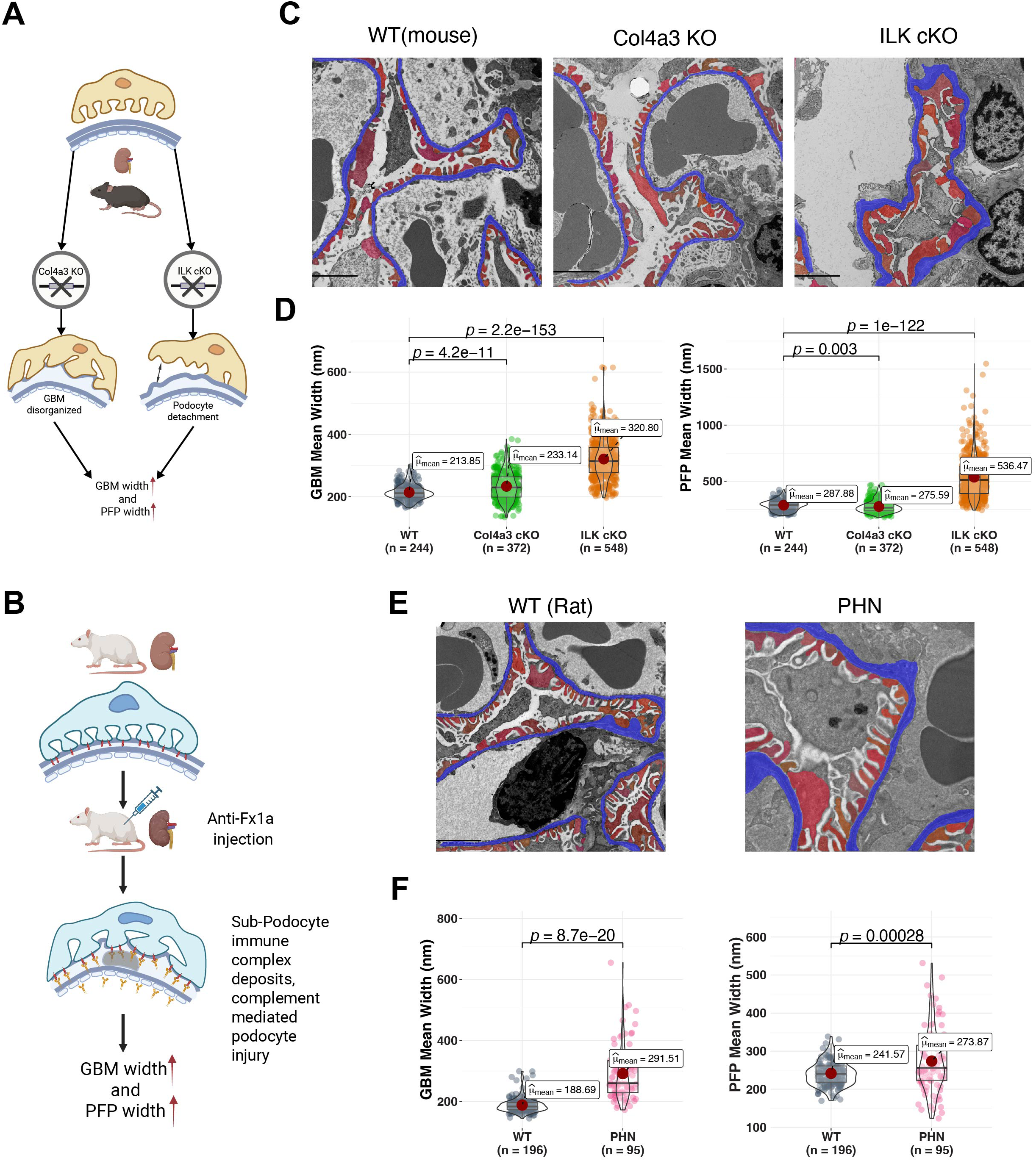
Application of TEAMKidney to three animal models of kidney disease. (A) Schematic illustrating two mouse models of glomerular disease: knocking out Col4a3 or specific podocyte ILK. (B) Schematic of a rat model of glomerular disease (PHN) induced by anti-Fx1a antibody injection. (C) Representative TEM images analyzed by the TEAMKidney model for WT, Col4a3 KO, and ILK cKO mice. (D) Violin plots showing quantitative measurements of PFP and GBM widths across WT, Col4a3 KO, and ILK cKO mice. (E) Representative TEAMKidney predicted images for WT and PHN rat samples. (F) Violin plots comparing PFP and GBM widths between WT and PHN groups.

We next examined the rat PHN model of immune-mediated podocyte injury^37^. In this model, rats were injected with anti-Fx1a antibody to induce the formation of immune complexes that deposit beneath podocytes and trigger complement-mediated podocyte injury, leading to progressive PFP effacement and GBM thickening (Fig. 3B, E). Automated measurements faithfully captured these ultrastructural changes: compared with WT animals, PHN rats showed a striking increase in GBM width (Fig. 3F). These differences are consistent with classical TEM-based characterizations of PHN, an animal model of human membranous nephropathy, confirming that our approach is sensitive to detecting pathological remodeling of the glomerular filtration barrier.

### Identifying Ultrastructural Remodeling in Human Kidney Diseases

TEM-based ultrastructural quantification is commonly used as diagnostic evidence in clinical renal pathology practice. To investigate the translational utility of our framework in human kidney disease states, we analyzed TEM images from patients with DKD and Fabry disease (Fig. 4A). DKD is characterized by hyperglycemia-induced hyperfiltration, GBM thickening, diffuse mesangial matrix expansion, leading to podocyte stress and progressive glomerular injury, PFP effacement, glomerular hypertrophy, and eventually podocyte loss and severe nodular glomerulosclerosis^49-51^. Fabry disease results from *GLA* mutations that cause accumulation of globotriaosylceramide (Gb3) in podocytes and subsequent structural damage, PFP effacement, podocyte detachment, and global or segmental glomerulosclerosis^52, 53^ (Fig. 4A). Representative TEM images predicted by our TEAMKidney model demonstrate marked distinct pathological features in both disease cohorts compared with normal controls (Fig. 4B).

**Fig. 4.**
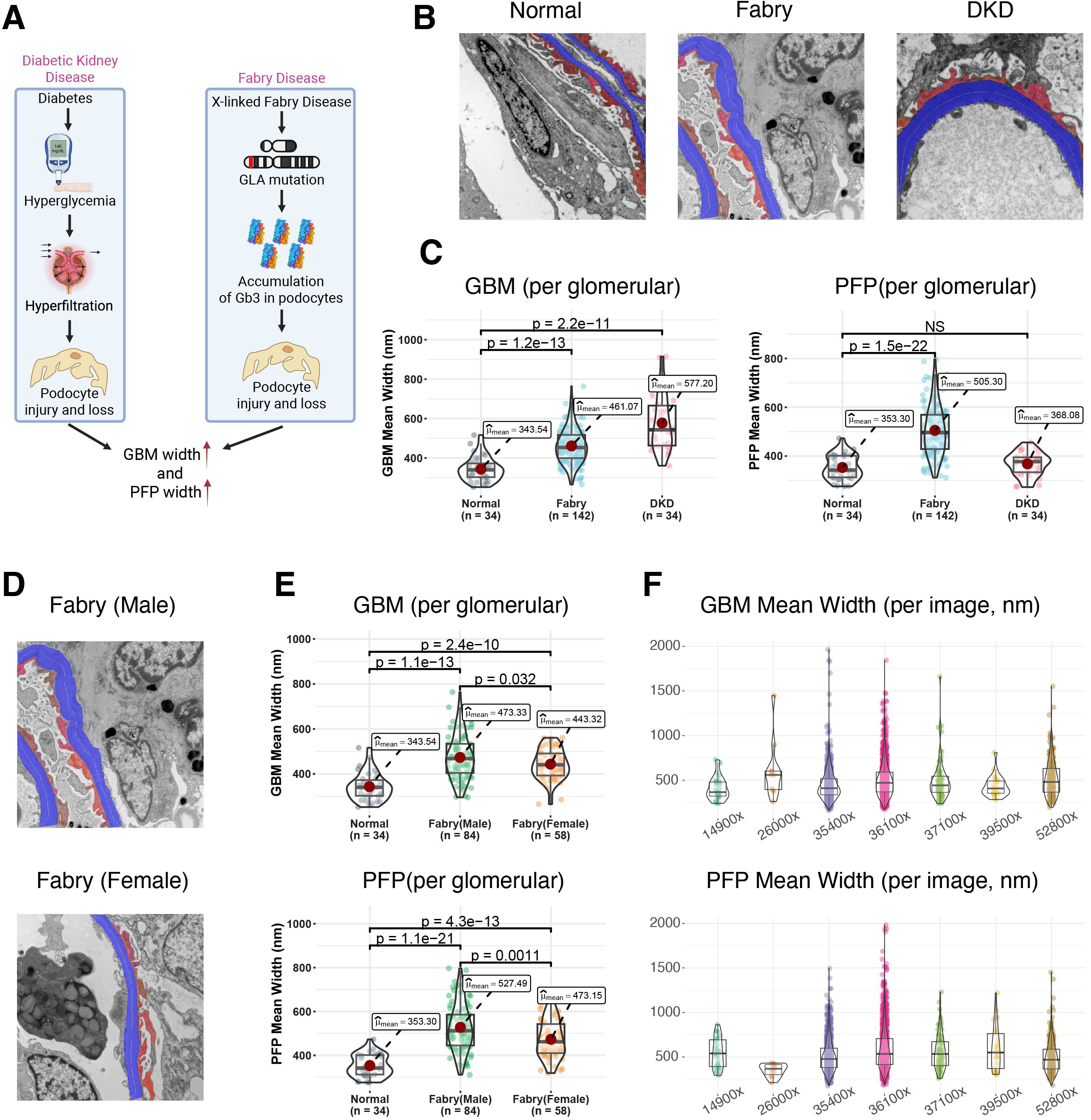
Application of TEAMKidney to human kidney diseases. (A) Schematic illustration of the pathogenesis of DKD and Fabry disease. (B) Representative clinical TEM images from normal, Fabry disease, and DKD patients were analyzed using the TEAMKidney model. (C) Violin plots showing the average GBM and PFP widths per glomerulus across normal, Fabry disease, and DKD samples. Each point represents the mean width of a single glomerulus. (D) Representative TEAMKidney predicted images from male and female Fabry disease patients. (E) Violin plots comparing GBM and PFP widths between male and female Fabry disease groups. (F) Violin plots illustrating the distributions of GBM and PFP widths across different magnifications in human kidney TEM images.

Quantitative measurements revealed disease-specific ultrastructural alterations (Fig. 4C). In DKD patients, GBM thickness was significantly increased. Notably, the mean PFP width in DKD showed only a slight increase compared with normal samples. The relatively preserved PFP width in our DKD cohort warrants further investigation, as it may reflect the disease stage of the included patients or capture an early phase where GBM pathology precedes overt podocyte architectural disruption. Longitudinal studies correlating PFP measurements with clinical disease staging would help clarify the temporal evolution of these ultrastructural changes^39, 54^. In contrast, patients with Fabry disease exhibited significant increases in both GBM thickness and PFP width, reflecting the more severe podocyte injury characteristic of lysosomal storage disorders^55, 56^.

Furthermore, sex-stratified analysis of Fabry patients revealed that both male and female patients exhibited substantial GBM and PFP widening (Fig. 4D, E), with more pronounced changes in Fabry males. These findings are consistent with the known pathology of Fabry disease, in which hemizygous males typically exhibit more severe phenotypes due to the X-linked inheritance pattern^57, 58^.

These results highlight the ability of our automated framework to detect clinically relevant and disease-specific ultrastructural alterations in human glomerulopathies. By accurately capturing the differential patterns of GBM thickening and PFP effacement in DKD and Fabry disease, the method demonstrates strong potential for application in both diagnostic pathology and clinical research, where reproducible and scalable measurements are critical for disease characterization, staging, patient stratification and management.

To assess robustness across imaging scales, we evaluated our method on TEM images acquired at multiple magnifications in the Fabry dataset. As shown in Fig. 4F, the predicted GBM and PFP mean widths remain consistent across different magnifications, despite substantial differences in image resolution. These results indicate that our method is stable and effective across imaging scales and disease conditions. Similar results for the DKD and normal datasets are shown in Supplementary Fig. 3.

### Comparison of PFP Quantification with a Benchmark Model

To better evaluate the performance, we benchmarked TEAMKidney against existing automated TEM quantification approaches using human, mouse, and rat kidney samples. As FPW-DL^31^ is the only method with publicly available code or model, and is designed exclusively to measure PFP width, the comparison was limited to PFP width quantification.

Considering FPW-DL was trained only on human samples, our comparison focused on human diseases. Even though all images were obtained from the FPW-DL study, TEAMKidney consistently outperformed FPW-DL across both normal and disease cohorts (Fig. 5A, B, Supplementary Fig. 4). Scatter plots show strong agreements between TEAMKidney and the ground truth for both PFP width and counts (Fig. 5A, B). In contrast, FPW-DL tended to overestimate PFP width and underestimate PFP counts, with this bias being more pronounced in DKD than in the normal cohort. Mean error of PFP width was reduced from 24.80 pixels (FPW-DL) to 4.46 pixels using our framework (Supplementary Fig. 4).

**Fig. 5.**
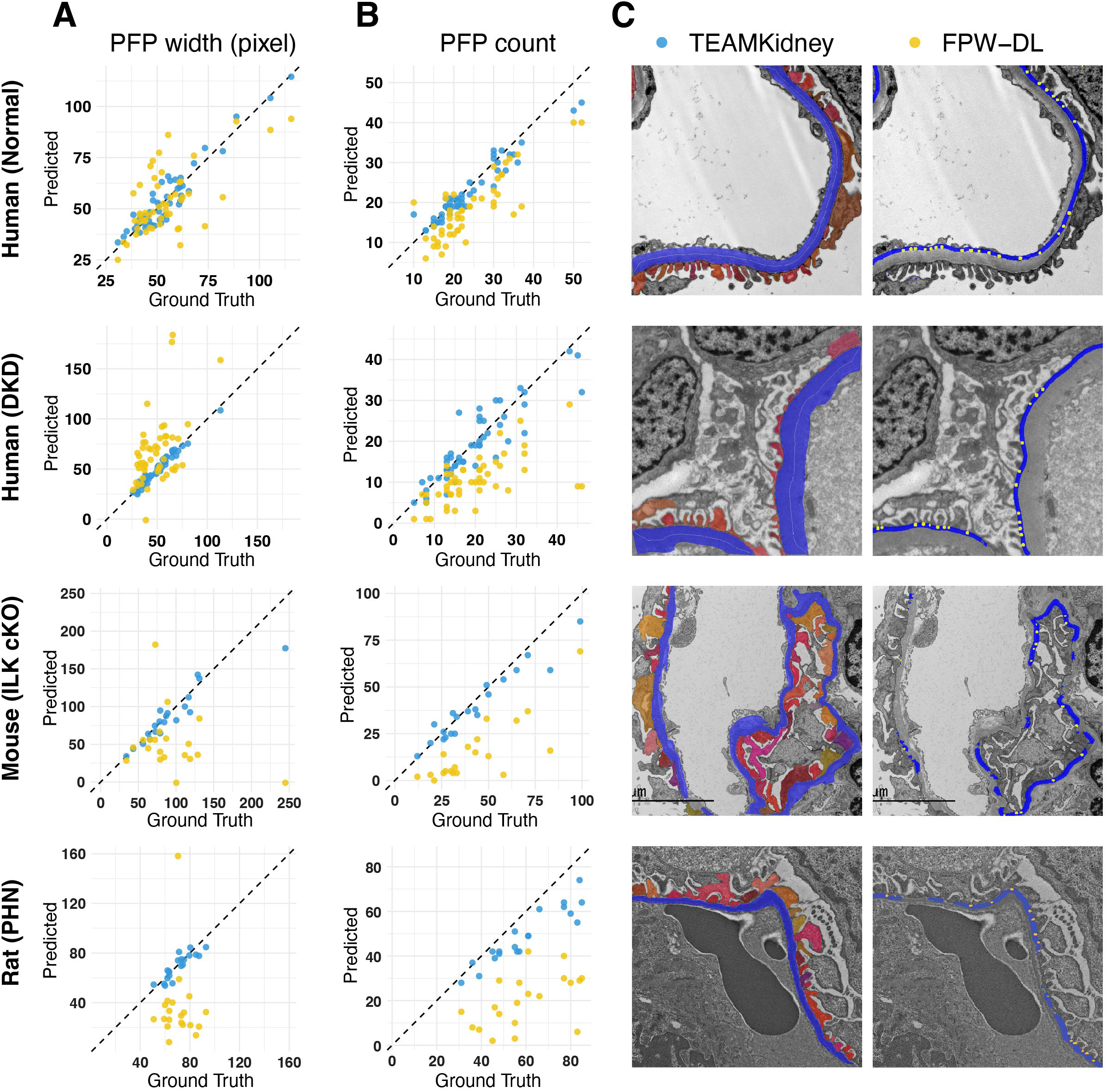
Comparison of PFP quantification between TEAMKidney and FPW-DL methods. Scatter plots with diagonal reference lines show the agreement between model predictions and ground truth measurements for (A) average PFP width and (B) PFP counts. (C) Representative TEM images with segmentation masks generated by TEAMKidney (left) and FPW-DL (right). From top to bottom: normal human kidney, human DKD, ILK cKO mouse model, and PHN rat model.

Further investigation revealed that PFP miscounting in FPW-DL arose from the use of slit diaphragm detection (yellow dots in Fig. 5C) rather than direct PFP detection. Under disease conditions or variable imaging quality, PFP effacement and overlap can obscure slit diaphragms, leading to overestimation of PFP width when slit diaphragms are missed, or preferential detection of relatively normal-appearing PFPs while pathologically altered regions are overlooked. Accurate ultrastructural quantification, therefore, critically depends on high-quality segmentation, which is the primary design goal of TEAMKidney. Representative TEM images show that our approach yields more continuous and anatomically faithful segmentation of podocyte FPs and the GBM compared with alternative methods (Fig. 5C), resulting in robust performance across both human disease states and animal models.

The performance gap between the two methods became more pronounced in animal sample (Fig. 5A, B), where FPW-DL, which was trained exclusively on human kidney data, showed substantially larger deviations from ground truth (Fig. 5A). This limitation stems from two primary factors: first, morphological differences between species, particularly the thinner GBM in rodents compared to humans, poses additional challenges for accurate structure identification; second, differences in imaging resolution and scale between our rodent datasets and FPW-DL’s training data further compromised its generalizability. In contrast, TEAMKidney maintained robust performance across species, demonstrating superior adaptability to diverse morphological features and imaging conditions.

### Agreement of TEAMKidney and Manual Measurements Across Species

Manually tracing and labeling each ultrastructural component is not feasible in practice. Instead, pathologists and biomedical scientists typically quantify ultrastructure by measuring multiple regions within an image to obtain representative values (see Methods). To assess discrepancies, we compared TEAMKidney measurements with manual measurements across normal, DKD, and Fabry disease (male and female) from human cohorts (Fig. 6) and animal models (Supplementary Fig. 5). For GBM widths, TEAMKidney quantification showed no significant differences from manual measurements in all groups (Fig. 6A, Supplementary Fig. 5), confirming that the framework preserves accuracy for this ultrastructural feature. Similarly, no significant differences were detected in PFP counts between TEAMKidney and manual counting across all groups (Fig. 6B, Supplementary Fig. 5), indicating that the framework maintains biological fidelity in enumerating PFP.

**Fig. 6.**
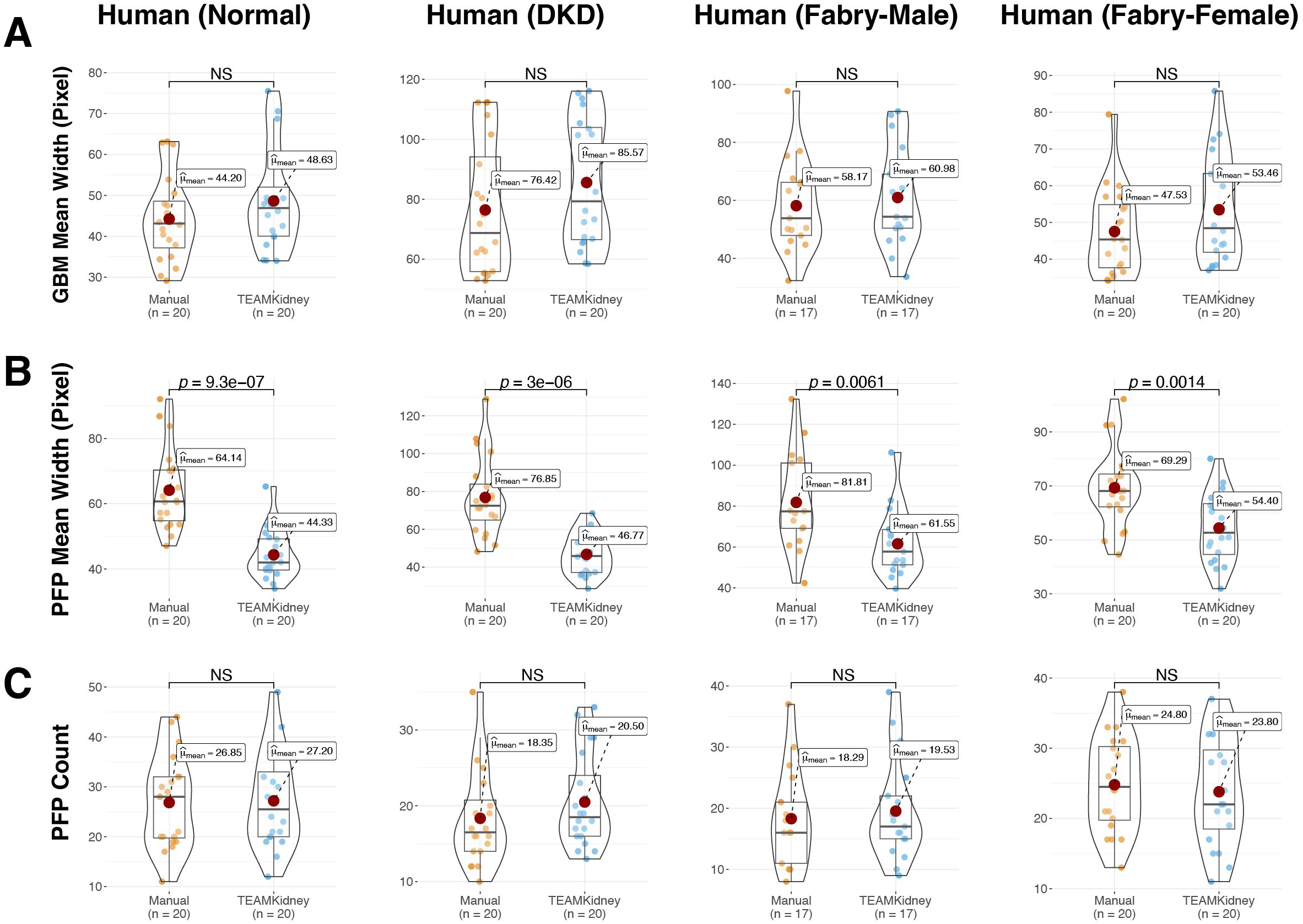
Evaluation of concordance between expert manual measurements and TEAMKidney predictions on human clinical TEM images. (A) GBM mean width. (B) PFP mean width. (C) PFP count. From left to right: Normal kidney samples, DKD samples, Fabry disease samples from male patients. (D) Fabry disease samples from female patients.

We observed a significant discrepancy in PFP width measurements between the two methods, with mean values from TEAMKidney consistently lower than those obtained by manual measurement. This difference likely arises from two factors. First, due to the irregular shape of PFPs, in the manual approach, PFP width is not measured directly but instead estimated by dividing the GBM length by the PFP count. GBM length is often overestimated, particularly in disease models with pronounced morphological alterations. Second, manual measurements include the slit diaphragm as part of the PFP, which inflates the estimated PFP width. In contrast, the TEAMKidney measurement strategy segments individual PFPs and slit diaphragm separately; therefore, the slit diaphragm is excluded from the PFP measurement, resulting in higher accuracy compared with the manual method (Fig. 6C, Supplementary Fig. 5).

Together, these results demonstrate strong concordance between TEAMKidney and manual measurements, establishing the framework as a reliable substitute for labor-intensive tracing. By accurately replicating expert-level quantification across both healthy and diseased human samples, the method enhances reproducibility and offers a scalable solution for ultrastructural kidney assessment.

### Comparison of Clinical Protocol and TEAMKidney for GBM and PFP Quantification

Although manual measurement is widely used in basic and clinical research, area selection is subject to individual judgment, leading to substantial inter-operator variability. In clinical settings, more stringent area-selection protocols are applied to improve reproducibility (see Methods). To contextualize the advantages of our automated approach, we compared it with the conventional diagnostic manual workflow commonly used for ultrastructural measurements of GBM and PFP widths (Fig. 7). Pathologists rely on ImageJ (or FIJI) based protocols, which require calibration with scale bars, grid overlays, and labor-intensive tracing of GBM thickness at multiple grid intersections, followed by foot process identification and marking of lateral edges (Fig. 7A). This procedure demands at least 200 measurements from three independent glomeruli per sample to achieve statistical robustness, making it time consuming and highly dependent on operator expertise.

**Fig. 7.**
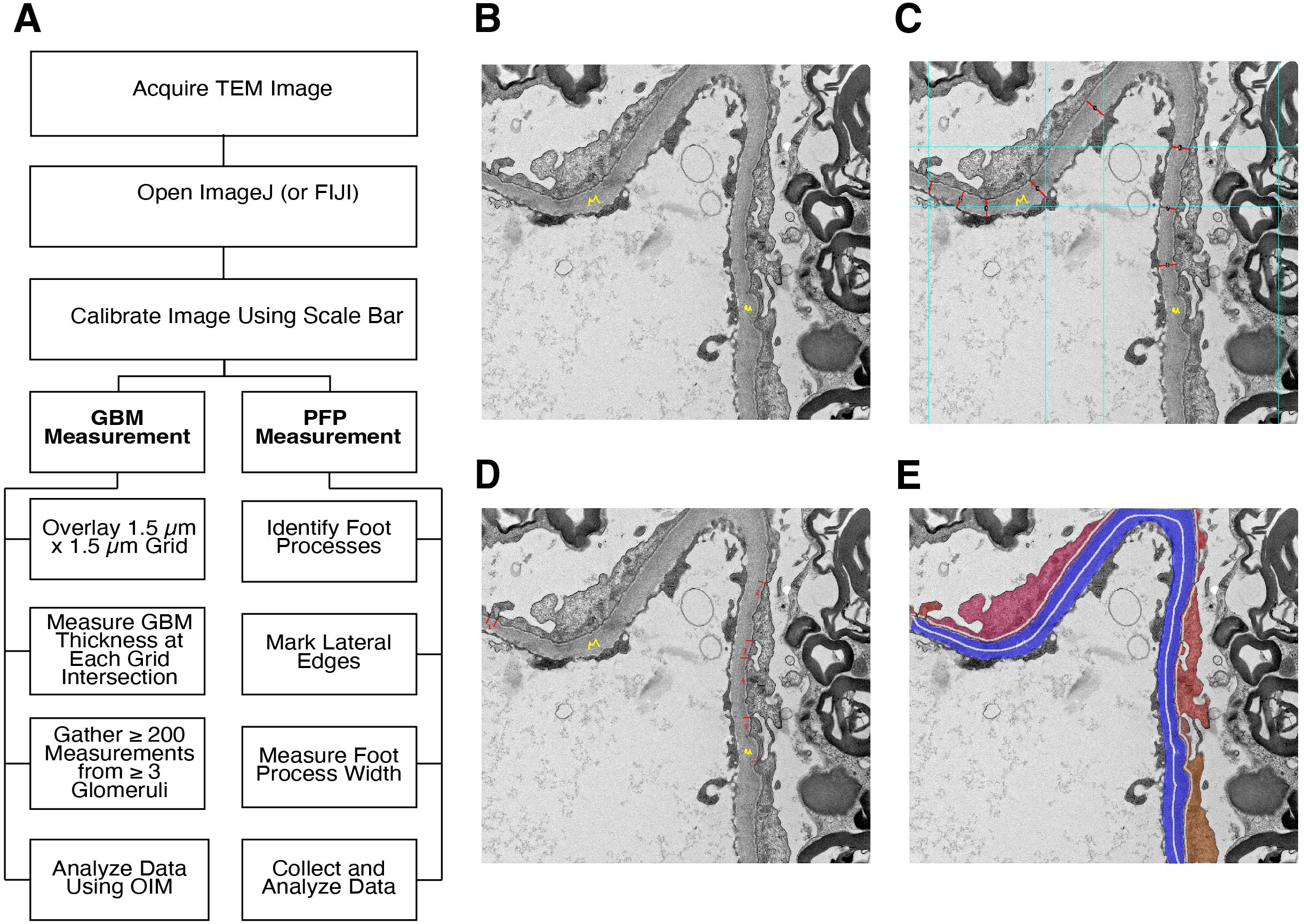
TEAMKidney aligned with clinical routine measurement protocols. (A) Schematic illustration of the clinical manual annotation process for GBM and PFP. Representative example showing (B) the original TEM image, (C) GBM measurement following the clinical manual procedure, (D) PFP counting following the clinical manual procedure, and (E) segmentation masks predicted by TEAMKidney.

Representative TEM images further illustrate the differences in measurement strategies (Fig. 7B-E). Manual quantification requires repetitive grid-based annotation and point-by-point width extraction (Fig. 7C and D), whereas our automated framework directly performs panoptic segmentation and downstream computational measurement of GBM and PFP widths (Fig. 7E). This not only eliminates observer variability but also enables high-throughput analysis of large datasets with consistent reproducibility.

Collectively, these comparisons underscore the limitations of traditional manual workflows, including their subjectivity, inefficiency, and limited scalability, and demonstrate that the proposed automated framework effectively overcomes these challenges, providing a practical and objective solution for ultrastructural kidney analysis.

## Discussion

In recent years, a global shortage of pathologists has been widely reported^59, 60^, affecting even large medical centers and resource-rich regions. The preparation and interpretation of tissue samples for TEM imaging require additional specialized training, particularly for renal pathologists, further exacerbating this workforce constraint. To address the urgent needs in nephrology research and clinical pathology, our team of experienced renal pathologists, nephrologists, biologists, and computational scientists designed and implemented TEAMKidney. Precise measurement of the GBM and PFP provides a much-needed solution to address labor-intensive and often subjective assessments of glomerular diseases. GBM and PFP are not only essential ultrastructure for diagnosing and monitoring chronic kidney diseases, but they also represent two distinct classes of TEM features. GBM is characterized by its continuous structure with a relatively regular shape and ambiguous boundaries throughout the image. In contrast, PFP consists of small, isolated structures with clear boundaries and irregular shapes. Accurate segmentation and quantification of these structures demonstrate the capability of the framework to extend beyond nephrology and renal pathology to general TEM image analysis across other fields.

High-quality labeled data is critical for training deep learning models, but manual annotation of TEM images is extremely time-consuming and requires specialized pathological expertise. To overcome this challenge, we adopted semi-automated pre-labeling strategies to accelerate the annotation workflow and implemented a multi-stage quality control process, ensuring that each label was independently reviewed and refined. These steps significantly reduced inter-operator variability and improved the overall reliability of the training data. Combined with a self-training strategy, our framework achieves high accuracy with only a few hundred finely annotated images, making it feasible to adapt to other tasks efficiently.

Previous TEM analysis methods have primarily focused on segmenting a single ultrastructural feature, often failing to generalize to other ultrastructures. Most studies have relied on U-Net based or semi-automated approaches (Table 2), which reduce the need for large amounts of training data but are well known for their limited generalizability to diverse morphologies. Because these methods typically use data from a single source and a single species, their practical applicability in animal disease modeling and clinical settings is further constrained. In addition to collecting and generating diverse datasets across multiple sources, species, and disease states, our specifically designed panoptic segmentation model offers two key advantages over directly applying generic computer vision models to this task. First, semantic segmentation provides spatially consistent, region-level guidance, reducing noisy or fragmented instance predictions during pseudo-labeling. Second, the trained HRNet model serves as the backbone for the panoptic segmentation network, enabling direct transfer and reuse of feature representations learned during semantic segmentation. This strategy not only accelerates model convergence but also improves feature consistency between semantic and panoptic tasks, thereby enhancing boundary precision and instance recognition. Overall, this hierarchical training strategy achieves greater stability and accuracy, particularly for densely packed ultrastructural components such as podocyte foot processes.

**Table 2.** Comparison Between Proposed Model and Other Published Methods.

| Method | Measurement of FP | Measurement of GBM | Label Dependency | Segmentation Target | DL model | Open Source | Year |
| --- | --- | --- | --- | --- | --- | --- | --- |
| Wu et al.<sup>22</sup> | No | Yes | N/A | N/A | N/A | No | 2010 |
| Yang et al.<sup>28</sup> | No | No | Supervised | GBMDD | U-Net | No | 2022 |
| Smerkous et al.<sup>29</sup> | Yes | No | Supervised | PGBMI, SD | U-Net | Yes | 2023 |
| Unnersjo-Jess et al.<sup>19</sup> | Yes | No | Supervised | PFP, SD | U-Net | No | 2023 |
| Lin et al.<sup>25</sup> | No | No | Self-supervised | GBM, PFP | GCLR | No | 2023 |
| Wang et al.<sup>23</sup> | No | Yes | Supervised | GBM | RADS-Net | No | 2024 |
| Laudon et al.<sup>24</sup> | Yes | Yes | Supervised | GBM | U-Net | Yes | 2025 |
| Yamashita et al.<sup>26</sup> | Only Effacement | Yes | Supervised | GBM, PFP | DeepLab V3 | No | 2025 |
| Zhang et al.<sup>27</sup> | No | Yes | Supervised | GBM, PFP | U-Net | No | 2025 |
| Ma et al.<sup>32</sup> | No | Yes | Supervised | GBM | YOLO+SAM | No | 2025 |
| **Our Method** | **Yes** | **Yes** | **Supervised** | **GBM, PFP** | **Glom2Mask** | **Yes** | **2025** |

TEM images inherently exhibit substantial variation in magnification and spatial resolution across different species, disease models, and experimental setups. Such differences often lead to inconsistencies in the visual scale and morphological representation of ultrastructural components, posing a major challenge for deep learning models trained on limited magnification ranges. To address this issue, we employed a series of strategies at multiple stages to enhance the model’s robustness to magnification variability. First, we collected TEM samples acquired at different magnifications across species and disease models to ensure comprehensive coverage of structural and contextual diversity. Second, we organized and standardized these multi-magnification samples by applying systematic cropping and rescaling operations to generate datasets representing diverse spatial scales. This step provided the model with balanced exposure to both high- and low-resolution features of the glomerular ultrastructure. Third, we incorporated random scaling, cropping, and other image augmentation techniques during model training to simulate continuous magnification changes and prevent overfitting to specific resolutions. Collectively, these strategies substantially improved the model’s ability to generalize across heterogeneous imaging conditions, yielding consistent segmentation quality and reliable quantitative measurements across a wide range of magnification differences.

Our proposed framework demonstrates high accuracy and reproducibility in segmenting and quantifying GBM and PFP, and generalizes effectively across species and disease states while accommodating substantial morphological and magnification variability. Nonetheless, there remains room for further improvement. Because TEM sections are two-dimensional representations of inherently three-dimensional structures, the spatial orientation of PFP can influence quantitative measurements, although this effect is unlikely to substantially alter overall conclusions when analyzed across whole images. To further enhance measurement accuracy, we are developing new methods that explicitly incorporate PFP orientation analysis. In parallel, we are extending the framework to include additional ultrastructural features relevant to other kidney diseases, such as immune deposits, slit diaphragms, and endothelial cells. As the first framework to achieve both segmentation and quantitative measurement with expert-level precision, TEAMKidney enables accurate characterization of glomerular morphological changes across diverse clinical conditions, ranging from normal kidneys and common diseases such as DKD to rare disorders such as Fabry disease. This work establishes a solid foundation for expanding the framework to quantify the full spectrum of glomerular diseases and to support future applications, including disease detection, classification, and automated report generation.

While open-source software is standard practice in computer science, it remains uncommon in the field of TEM image analysis. With the exception of FPW-DL, all other existing studies have not released their code or trained models, making independent evaluation impossible and significantly limiting the broader adoption and impact of these methods. We greatly respect the FPW-DL authors for openly sharing their code, models, and data, and for their prompt and helpful responses to our inquiries. We strongly believe that making our own code and models publicly available will foster community-wide experimentation, accelerate methodological advances, and ultimately benefit patient care. This openness also reflects our commitment to transparency and confidence in the robustness of our approach.

## Methods

### Sample Preparation and Image Acquisition

#### ILK podocyte-specific knockout (ILK cKO) and Col4a3 knockout (Col4a3 KO) models

The ILKcKO mice were generated by crossing ILK^flox/flox^ mice with Podocin-Cre^+^ mice^36^. Col4a3 KO mice were purchased from the Jackson Laboratory (RRID:IMSR_JAX:002908). The littermate WT mice were used as healthy controls. Mice were sacrificed at 16 weeks of age, and the kidneys were dissected. The fresh kidney samples were fixed in Karnovsky’s fixative and processed according to the published protocol^4^. In brief, small pieces of kidney samples (<1 mm thick) were fixed in buffered 2.5% glutaraldehyde. Kidney specimens were then post-fixed in 1% osmium tetroxide (Electron Microscopy Sciences), dehydrated, embedded in propylene oxide/Epon resin, and sectioned (60 to 90 nm thick) using the ultramicrotome. Collected sections were post-stained with uranyl acetate and lead citrate (Electron Microscopy Sciences), according to the published protocol^4^. Images were obtained using the Jeol JEM-1011 transmission electron microscope (JEOL USA Inc.) equipped with an Erlangshen ES100W digital camera (Gatan, Pleasanton, CA). The operator was blinded as to the genotype and the injury status of the animals at the time of image collection and analyses^16^.

#### Rat PHN injury model

A modified PHN injury model was induced in Lewis rats (strain code 004; Charles River Laboratories, Wilmington, MA) by one dose of i.v. injection [1.5 to 2 mL/kg sheep anti-rat Fx1A serum (PTX-002S; Probetex, San Antonio, TX)], according to Probetex and published protocol^61^. Rats were euthanized at day 16 after anti-rat Fx1A serum injection, and kidneys were collected and fixed in Karnovsky’s fixative for TEM analysis. Sterile 0.9% sodium chloride (normal saline) was used in control rats. A total of six animals in each experimental group were studied. Kidney samples were prepared, and TEM imaging was performed as described above.

#### Human diseases

TEM images used in this study were obtained from an open-source dataset described in a recent publication^31^. The dataset includes kidney biopsy samples from patients with Fabry disease, type 2 diabetes, and minimal change disease, as well as healthy controls from living kidney transplant donors. All biopsies were processed at eight EM laboratories following standardized protocols. Ultrathin glutaraldehyde fixed sections were stained with toluidine blue for glomerular identification, and high magnification transmission EM images (original magnification ×30,000) were captured using a systematic uniform random sampling strategy.

### Label Generation, Manual Measurement, and Image Prepressing

#### Ground Truth Label

All TEM images were divided into training, validation, and testing datasets as summarized in Table 1. The training/validation and testing sets were intentionally sourced from different cohorts, encompassing distinct species and disease types, to ensure that the testing dataset could robustly evaluate the generalization capability of our algorithm. For the training and validation sets, regions corresponding to the GBM and PFP were manually annotated using LabelMe software^62^. The annotations were performed by trained researchers using high-resolution TEM images and subsequently reviewed by renal pathologists to ensure consistency and structural accuracy. This manual labeling process served as the foundation for model training and validation, while the independently collected testing data were reserved for assessing model performance across unseen imaging and disease conditions.

#### Manual measurement for GBM and FP widths

Manual podocyte foot process width was traced and measured on ×12,000 high magnification TEM images using ImageJ software version 1.47v, according to published methods^16^. Briefly, the total length of the GBM spanning the area of the slit diaphragms was measured in each image using ImageJ, according to published methods^17, 63,64^. The results were then pasted on an Excel sheet and adjusted for the scale bar. After that, all slit diaphragms covering the GBM were counted, and the PFP width was calculated using the total length of GBM divided by the total number of slit diaphragms in each image. The GBM width was measured as the distance from the podocyte membrane to the endothelial membrane. Each image was randomly measured in 20 areas around the GBM, and the harmonic mean was calculated for each image.

#### Preparing training and testing datasets for the deep learning framework

Images were normalized by mean-std normalization and tiled into [size, e.g., 1024×1024, 2624*2624] with 50% overlap. Scale bars and metadata were parsed to record the pixel-to-nm factor (e.g., 4.122 nm/pixel for mice, 3.88 nm/pixel for rats) used for unit conversion. To avoid leakage, splits were patient/animal-level stratified into train/validate with an 80/20 ratio while preserving class balance per species and disease model. All preprocessing scripts are provided (see Code availability).

#### Hybrid Data Preparation

To improve the model’s adaptability across images of different resolutions, we adopted a stage-wise data preparation and training strategy. High-resolution TEM images were first used to pretrain the backbone network, enabling the model to learn fine-grained structural features of GBM and PFP. Subsequently, lower resolution images were used for fine-tuning to encourage scale-invariant feature learning and improve cross-magnification generalization. During both training stages, various data augmentation techniques were applied, including random scaling, random cropping, horizontal and vertical flipping, and random adjustments of brightness and contrast. These augmentations helped prevent overfitting and further enhanced the model’s robustness to variations in image scale and quality.

#### Ultrastructure Quantification Based on Clinical Protocols

GBM thickness and PFP width were measured using TEM images analyzed in ImageJ (FIJI). Images were calibrated using reference scale bars to ensure accurate morphometric analysis. For GBM thickness measurement, a standardized 1.5 µm × 1.5 µm grid was superimposed on micrographs for systematic sampling. At each grid intersection, perpendicular lines were drawn from the endothelial cell membrane to the outer margin of the lamina rara externa beneath epithelial foot processes. Measurements were taken only in well-preserved regions, excluding artifacts due to sectioning angle, folding, or detachment. The Orthogonal Intercept Method (OIM) was applied to calculate mean GBM thickness. For PFP width measurement, the lateral edges of individual PFPs were manually identified, and width was measured as the linear distance between adjacent borders. Only clearly defined and intact PFPs were included in the analysis.

### Glom2Mask: TEM Image Panoptic Segmentation Model Architecture

Our proposed Glom2Mask model is designed based on the Mask2Former panoptic segmentation framework, enabling comprehensive segmentation of GBM and PFP in TEM images. Given the distinct structural characteristics of these components, we categorize GBM as a "stuff" class for semantic segmentation and PFP as a "things" class for instance segmentation. This formulation ensures precise differentiation between the continuous structure of the GBM and the discrete PFP instances within the glomerular filtration barrier.

To enhance segmentation accuracy, we integrate HRNet as the backbone for high-resolution feature extraction. HRNet is well-suited for this task due to its ability to preserve spatial details throughout the network, which is essential for capturing fine structural differences in TEM images. Unlike traditional backbones that downsample feature maps at early stages, HRNet maintains high-resolution representations, allowing for better delineation of GBM boundaries and accurate identification of individual PFP structures.

Our model follows a transformer-based approach, where multi-scale features extracted by HRNet are processed through multi-head attention mechanisms to generate high-quality segmentation masks. The network outputs a set of learned queries, each corresponding to a GBM region or a PFP instance, enabling robust panoptic segmentation across diverse TEM datasets, including mouse, rat, and human kidney samples.

By leveraging HRNet for high-resolution feature extraction and Mask2Former for unified panoptic segmentation, our model effectively segments GBM and PFP structures, providing the necessary groundwork for precise automated width measurement in kidney disease analysis.

### Model Training for the Deep Learning Framework

Our training process consists of two stages: a self-training-based semantic segmentation stage and a panoptic segmentation stage using Glom2Mask. The first stage aims to generate additional pseudo-labeled data while simultaneously learning a high-resolution backbone network. In the second stage, we utilize the HRNet trained in the first stage as the backbone of a modified Mask2Former model for panoptic segmentation, thereby improving both training efficiency and segmentation accuracy.

#### Self-Training for Semantic Segmentation

In the first stage, we employ a self-supervised learning approach to HRNet-based semantic segmentation.

For cases with a limited number of annotated images (e.g., 19, 48, or 64 labeled samples), we employed a self-training strategy to improve segmentation performance. Self-training is a semi-supervised learning technique that utilizes both labeled and unlabeled data. Initially, the model is trained using only the labeled dataset to capture fundamental feature representations. After this initial training, the trained model generates pseudo labels for the unlabeled data. These pseudo-labeled samples, combined with the original labeled dataset, are then used to retrain the model iteratively, progressively refining its segmentation accuracy and generalization capabilities. The model performance was evaluated using the mean Dice score. For each manual segmentation mask, per category, was compared to the prediction mask of the model. Dice score was calculated by,

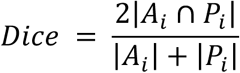

Where *A_i_* as the ground truth segmentation result for class *i, P_i_* as the predicted segmentation result for class *i*. Dice score values range from 0 to 1, where 0 represents no pixel overlap, and 1 represents perfect pixel overlap between the two masks.

During training, we applied a series of data augmentation techniques to improve model robustness, including random scaling within a range of 0.5-2.0, random rotation within ±10°, horizontal and vertical flipping with a probability of 0.5 each, random cropping to a size of 1024×1024 pixels, and random intensity jitter with brightness and contrast factors between 0.8 and 1.2. The model was optimized using the AdamW optimizer with a learning rate of 6×10^-5^ and a weight decay of 0.05. A polynomial learning rate decay schedule with a linear warm-up phase of 1,500 iterations was employed. Training was conducted with a batch size of 16 for a total of 80,000 iterations. To address class imbalance, a combination of cross-entropy loss and focal loss (γ = 2.0, α = 0.25) was used.

*Panoptic Segmentation with Glom2Mask*.

In the second stage, the training setup follows the same training validation split as in the first step, ensuring consistency in dataset partitioning. Data augmentation techniques, including random scaling, cropping, rotation, flipping, and contrast adjustments, are applied during training to enhance generalization similar to step 1. The model was optimized using the AdamW optimizer with a learning rate of 1 × 10^-5^ and a weight decay of 0.05 for a total of 160,000 iterations. The training objective combined three loss functions, including classification cross entropy, mask cross entropy, and Dice loss, with relative weights of 2.0: 5.0: 1.0. Early stopping and checkpoint selection were determined based on the PQ metric evaluated on the validation set.

The loss functions for training are as follows: Segmentation classification head: Cross-entropy loss, ensuring accurate categorical predictions for GBM and PFP. Mask prediction head: A combination of cross-entropy loss and Dice loss, improving both pixel-wise accuracy and mask shape consistency. To quantitatively assess model performance, we employ the following panoptic segmentation metrics:

- Panoptic Quality (PQ): Measures the overall segmentation accuracy, balancing segmentation precision (true positive detection) and quality (IoU based mask accuracy). Defined as:

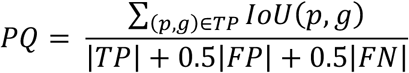

where TP (true positives) are correctly matched masks, FP (false positives) are extra masks, and FN (false negatives) are missed masks.
- Segmentation Quality (SQ): Measures the average mask accuracy of correctly predicted objects:

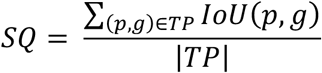
- Recognition Quality (RQ): Measures the correctness of mask instance association and is computed as:

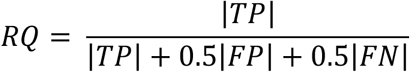

### Training Environment and Reproducibility

All experiments were run on Ubuntu 22.04 with Python 3.9, PyTorch 2.1.0, CUDA 12.1, cuDNN 8.9, MMDetection 2.0.1, and MMCV. Hardware:1*NVIDIA A100-80G GPU, Intel Xeon Gold 6338 CPU (40 cores). Deterministic behavior was enforced using a fixed seed (seed=42) and disabling nondeterministic cuDNN ops. Full environmental files, exact config YAMLs, and trained checkpoints are provided (see Code availability).

### Post Segmentation Processing and Quantification

#### GBM Width Measurement

The second stage of our proposed method consists of a Python image processing algorithm that takes input of the GBM and PFP segmentation tiles predicted by the deep learning model to measure the mean GBM width and PFP width. We will describe the GBM width measurement first here. The mean GBM width for each tile is then calculated as its total GBM area divided by its total GBM central length according to the following equation

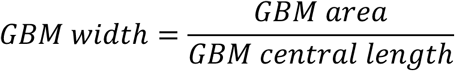

Where the area of the GBM is straightforward to compute, it involves directly counting the number of pixels in a section of the GBM. The process for calculating the length of the GBM is as follows: First, we obtain the skeleton of the GBM using the skeletonize method with the Python package scikit-image. The general skeleton obtained through the skeletonize method, in addition to the central length of the GBM, also includes many small branching points (Fig. 1J). To eliminate these branching points, we construct an undirected cyclic graph of the GBM skeleton using the Python package NetworkX. Here, there are two scenarios: if the GBM forms a loop structure, we iteratively prune nodes with an in-degree of one from the undirected cyclic graph of the GBM skeleton until no nodes with an in-degree of one remain. We then calculate the perimeter of the resulting graph to determine the central length of the GBM; if the GBM forms a strip-like structure, we calculate the shortest distance between any two endpoints and identify the two endpoints with the greatest distance as the central length of the GBM. Lastly, the specimen mean GBM width was converted from pixels to distance with a conversion factor derived from the TEM image scale bar, for example, 3.88 nm/pixel for rat model data.

#### PFP Width Measurement

Following this, the PFP width calculation is more direct. Each PFP instance is represented by a unique integer starting from 1, with the background assigned a value of 0.

Next, we use morphological operations in OpenCV to dilate the GBM, enabling us to obtain its contour. In this processed image, the background remains coded as 0, and the contour positions are coded as 1.

Finally, we calculate the width of each PFP by multiplying the PFP instance matrix by the matrix of the dilated GBM contours. This operation yields the width by computing the pixel length of PFPs. Similarly, the specimen PFP widths were converted from pixels to distance with a conversion factor derived from the TEM image scale bar, for example, 3.88 nm/pixel for rat model data.

## Author contributions

C.Z., W.L., A.Z., and X.F. conceived the study. A.Z. developed the deep learning model. W.L., X.F., S.R.V., S.P.B., H.Y., and H.C. acquired the images from animal models. X.F., A.Z., W.T., J.J., L.D., and E.O. labeled, curated, and measured the images. A.Z., W.T., J.J., and C.Z. performed the data analysis. F.R.M. and J.M.H. provided clinical consultation and measurements. A.Z., C.Z., X.F., W.T., and J.J. generated the figures and wrote the original draft. C.Z., W.L., X.F., and A.Z. supervised the work and revised the final manuscript. All authors reviewed and approved the final manuscript.

## Data availability

The datasets supporting this study’s findings are publicly accessible and include three infrared and visible image datasets: human disease dataset (https://github.com/najafian-lab/fpw-dl-v1), mice and rat disease model dataset (https://github.com/CZCBLab/TEAMKidney)

## Code availability

The code is available at https://github.com/CZCBLab/TEAMKidney.

## Acknowledgments

We thank Casey Ritenour, Cheryl Tyszkiewicz, Seo-Kyoung Hwang, Chang-Ning Liu, and Lindsay Tomlinson of Pfizer Global Pathology and Comparative Medicine and Drug Safety R&D for help with rat PHN models. We thank Richa Sharma, Sudhir Kumar, and Boston University’s Animal Science Center for their help in generating Col4a3 and ILK knockout mice. This work is supported by research grants from NIH/NIDDK grants R21DK143406 (CZ, WL), R01DK133940 (WL), R01DK078226 (WL), and a DOD grant E01 HT9425-23-1-1058 (WL). This work is also supported by BIT-ARC and B4D-ARC grants from Boston University Evans Center (WL, CZ, JMH), a grant from Pfizer Centers for Therapeutic Innovation (WL), a BU CTSI Integrated Pilot Grant funded by the Department of Medicine (CZ, WL), and the Boston University Undergraduate Research Opportunities Program (JJ). This publication is also supported in part by the National Center for Advancing Translational Sciences, National Institutes of Health, through BU-CTSI Grant Number 1UL1TR001430. Its contents are solely the authors’ responsibility and do not necessarily represent the official views of the NIH. The funders had no role in study design, data collection and analysis, decision to publish, or preparation of the manuscript

