## Supplementary material for "Deep Learning Enables Automated Segmentation and Quantification of Ultrastructure from Transmission Electron Microscopy Images"

**SUPPLEMENTARY MATERIAL TABLE OF CONTENTS**

1. **Supplementary Methods.**
2. **Supplementary Fig. 1. Representative TEM images from Mouse and Human samples segmented by the three backbone networks for** **qualitative comparison.**
3. **Supplementary Fig. 2. Evaluation of Glom2Mask robustness across different input image resolutions.**
4. **Supplementary Fig. 3. Evaluation of TEAMKidney robustness across human kidney diseases samples with different magnifications.**
5. **Supplementary Fig. 4.** **Comparison of FP quantification between TEAMKidney and FPW-DL methods in additional models.**
6. **Supplementary Fig. 5.** **Evaluation of concordance between expert manual measurements and TEAMKidney predictions on human clinical TEM images.**
7. **Supplementary Table 1.** **Performance comparison between the proposed Glom2Mask model and the benchmark Mask2Former for panoptic segmentation.**

**Supplementary Methods**

**Dataset Construction and Model Training Pipeline**

As Figure 2A shown, the pipeline consists of three major stages. In the first stage (Self-Training & Semantic Segmentation), the initial HRNet-FCN model (Step 1) is trained on a small, accurately labeled dataset (DS1, consisting of 19 mouse images) and used to generate predictions on the unlabeled dataset (DS2, comprising 96 mouse, 540 rat, and 470 human images). Following this initial prediction, 24 mouse, 107 rat, and 93 human images are selected from DS2 and manually annotated with fine-grained labels to form a validation subset, which serves both as a benchmark for evaluating semantic segmentation performance across self-training iterations and as the evaluation reference for the subsequent panoptic segmentation model. This validation subset is strictly excluded from all subsequent self-training iterations and does not participate in any further pseudo-label generation or model training. Thereafter, the model performs multiple rounds of iterative training exclusively on the remaining non-validation images in DS2 (72 mouse, 433 rat, and 377 human images): in each round, the model predicts on these unlabeled images to generate a pseudo-labeled dataset (DS3, with the same composition as the remaining data), and the pseudo-labels are then used to retrain the model, progressively improving label quality through repeated cycles. The final pseudo-labeled DS3 undergoes manual curation to produce a high-quality, accurately labeled dataset DS4.

In the second stage (Panoptic Segmentation), the accurately labeled dataset DS4 (comprising 115 mouse, 540 rat, and 470 human images) is split into training and validation sets following the same partition strategy as in the first stage, and is used to train the Glom2Mask model (Step 3). The HRNet backbone obtained through self-training in Step 1 is adopted as the backbone of the Glom2Mask panoptic segmentation model, enabling the model to leverage the feature representations learned during self-training. Model performance is evaluated on the validation subset, which did not participate in the first-stage training.

In the third stage (Performance Evaluation), the trained Glom2Mask model is applied to the evaluation dataset (DS5, comprising 328 mouse, 113 rat, and 10,099 human images), and model outputs are assessed across two dimensions: GBM/PFP width measurements and downstream biological analysis.

**GBM Skeleton Extraction**

In our initial implementation, we obtained a one-pixel-wide skeleton of the segmented GBM by directly skeletonizing the binary GBM mask. However, for thin band-like GBM regions this naïve skeletonization frequently produced numerous small side branches. These spurious branches artificially increased the effective GBM length and led to an overestimation of the GBM cross-section, thereby biasing the downstream computation of mean GBM width. To address this issue, we developed an improved skeleton extraction strategy that explicitly suppresses side branches and retains only a single, biologically meaningful centerline.

Concretely, we first detect all contours in the binary GBM mask using OpenCV and sort them by contour length. The ratio between the lengths of the two longest contours (ctLen_1_ and ctLen_2_) is used to distinguish approximately ring-like GBM profiles (0.5 < ctLen_1_/ctLen_2_ < 2) from band-like profiles (otherwise). For ring-like profiles, we retain only the two main contours and fill all smaller contours; for band-like profiles, we retain only the largest contour and fill all remaining ones. The resulting cleaned GBM mask is then skeletonized. Very small skeletons (total number of skeleton pixels < 500) are discarded as unreliable detections.

For the remaining skeletons, we construct a graph-based representation to prune branches and extract the dominant centerline. We identify skeleton endpoints and bifurcation points and treat them as graph vertices. Each skeleton branch connecting a pair of vertices is traced and stored as an edge whose weight equals its pixel length. Using this weighted graph, we apply Dijkstra’s algorithm to compute the shortest-path distance between all vertex pairs and select the pair with the maximal geodesic distance. We then reconstruct the corresponding sequence of skeleton pixels by concatenating the stored branch segments along this longest path. The resulting centerline skeleton contains only the main GBM trajectory with side branches removed, providing a more accurate representation of the GBM path for subsequent GBM width estimation.

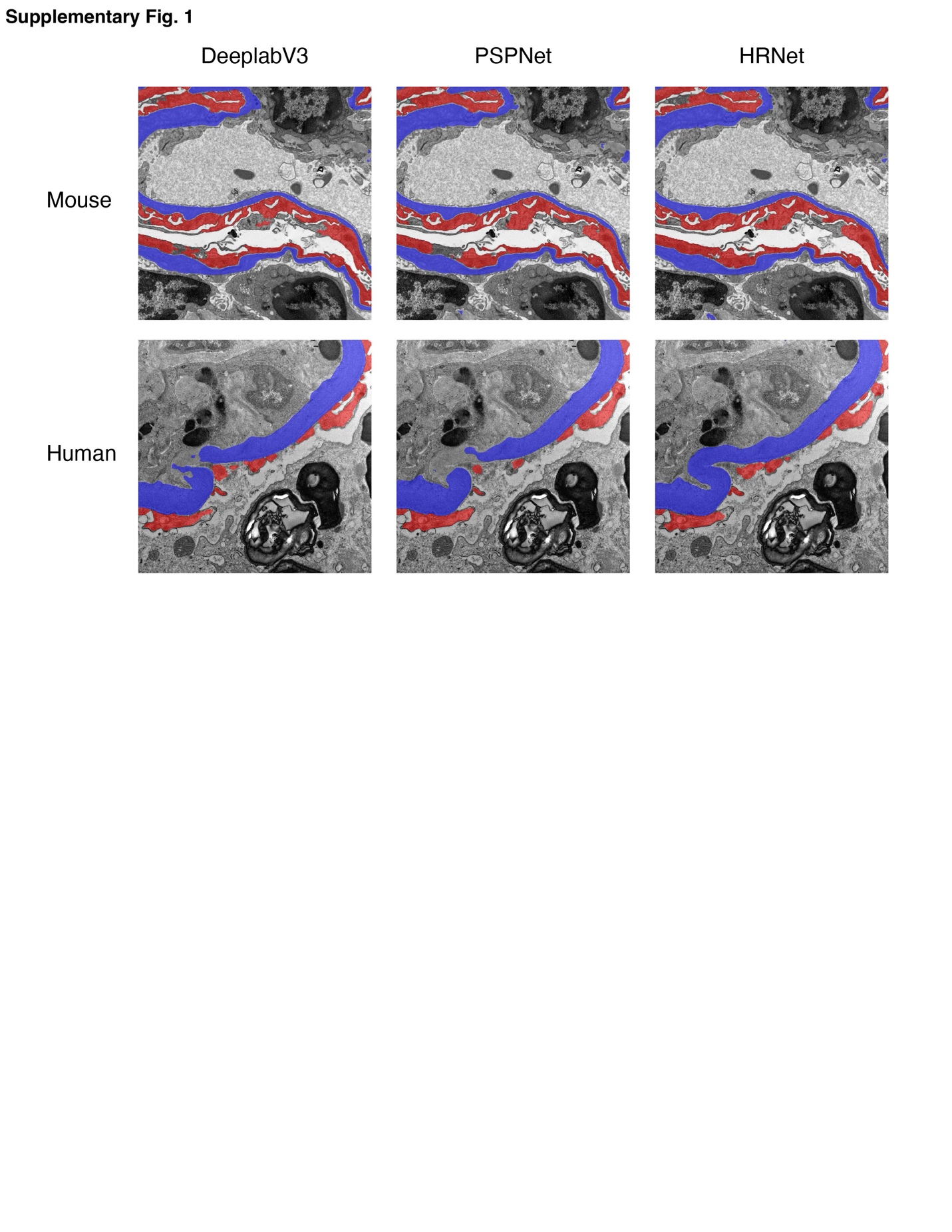

**Supplementary Fig. 1. Representative TEM images from Mouse and Human samples segmented by the three backbone networks for qualitative comparison.**

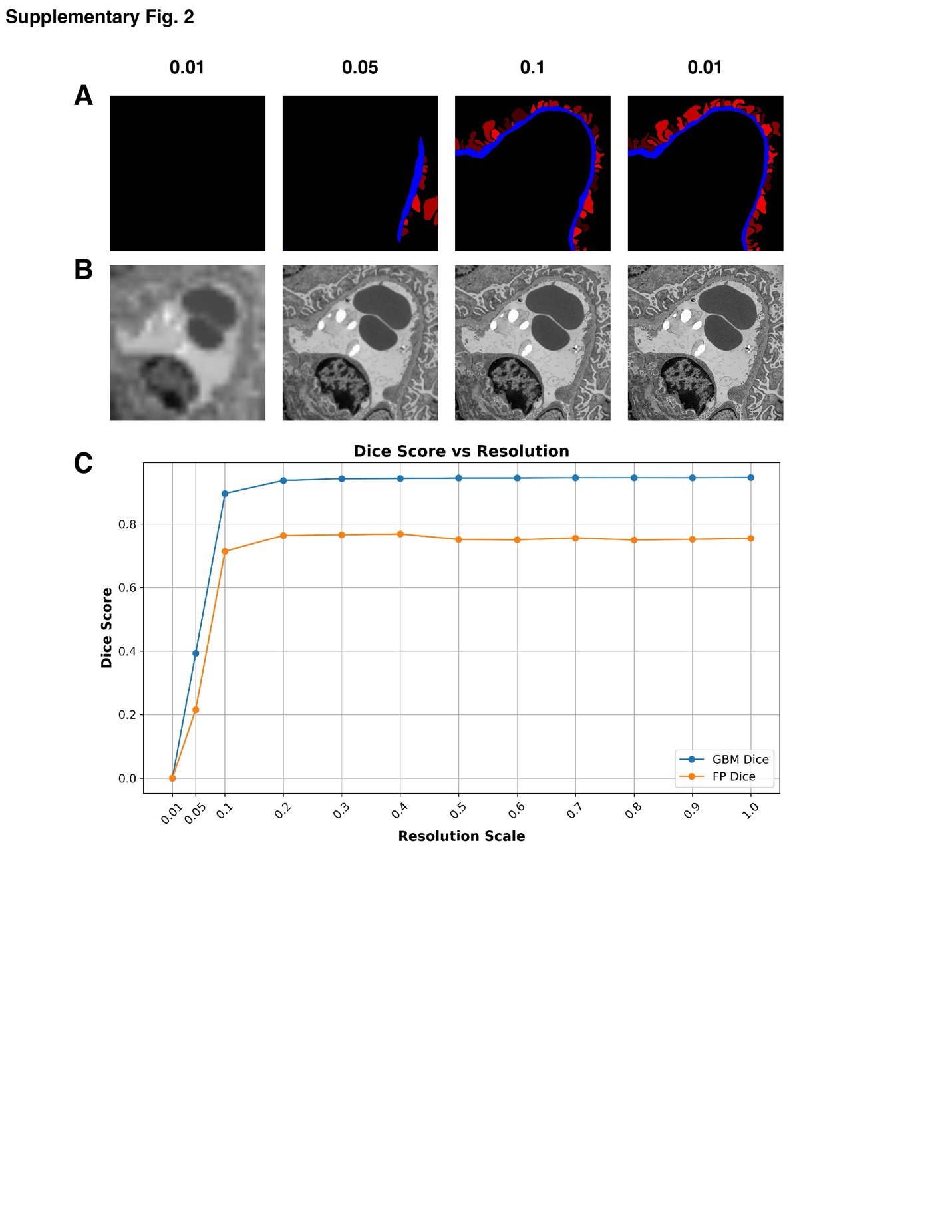

**Supplementary Fig. 2. Evaluation of Glom2Mask robustness across different input image resolutions.** (A) Representative segmentation results of GBM (blue) and FP (red) produced by Glom2Mask at different resolution scales, where values from 0.01 to 1 indicate the downsampling ratio relative to the original TEM images. Progressive improvement in boundary delineation is observed with increasing resolution. (B) Corresponding TEM input images at each downsampling ratio, illustrating substantial differences in image detail and ultrastructural visibility across scales. (C) Quantitative evaluation of segmentation performance as a function of the downsampling ratio, measured by Dice scores for GBM and FP. Segmentation accuracy improves rapidly with increasing resolution and remains stable over a wide range of downsampling ratios, demonstrating robustness of the proposed method to resolution variation.

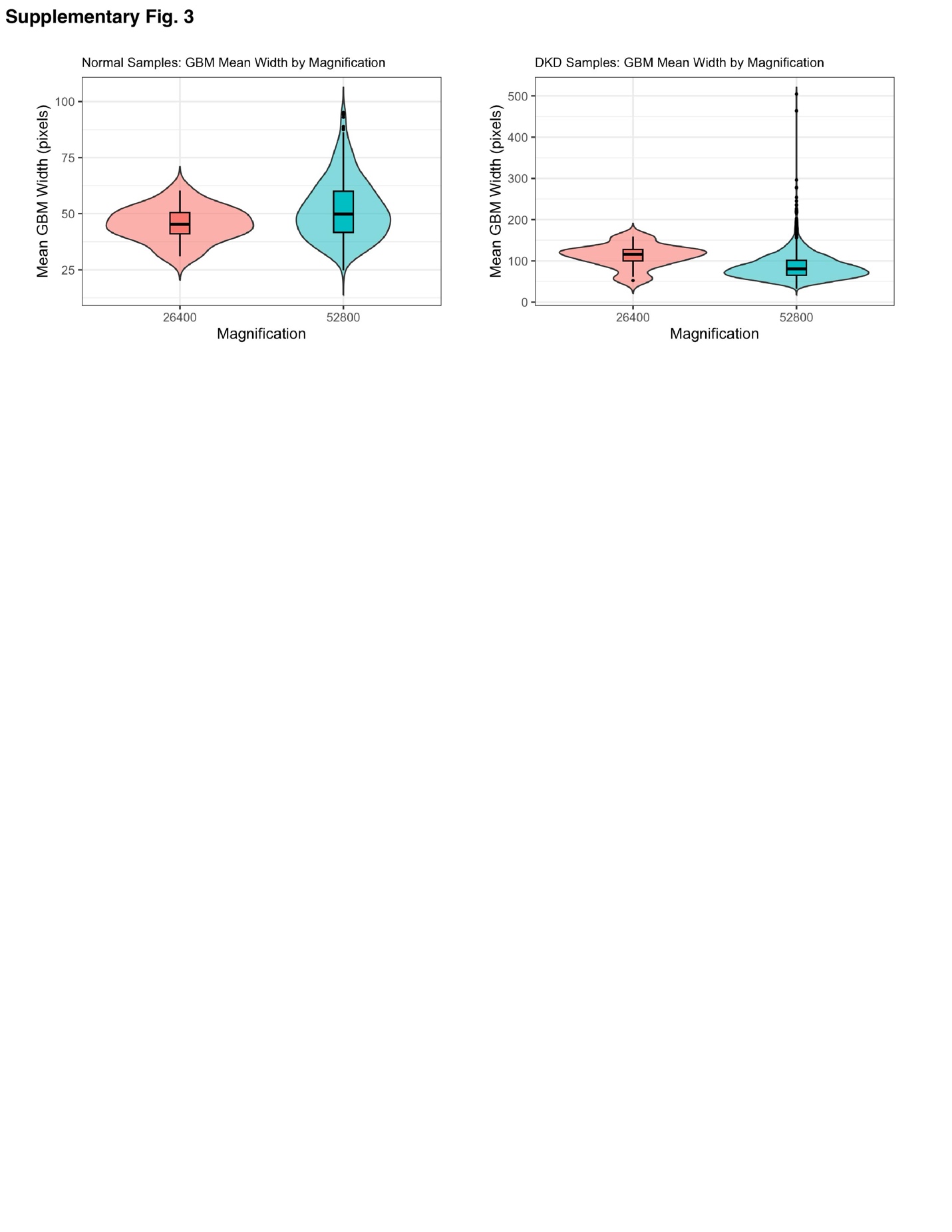

**Supplementary Fig. 3. Evaluation of TEAMKidney robustness across human kidney diseases samples with different magnifications.** Violin plots illustrating the distributions of GBM across different magnifications in TEM images from normal human samples and DKD patients.

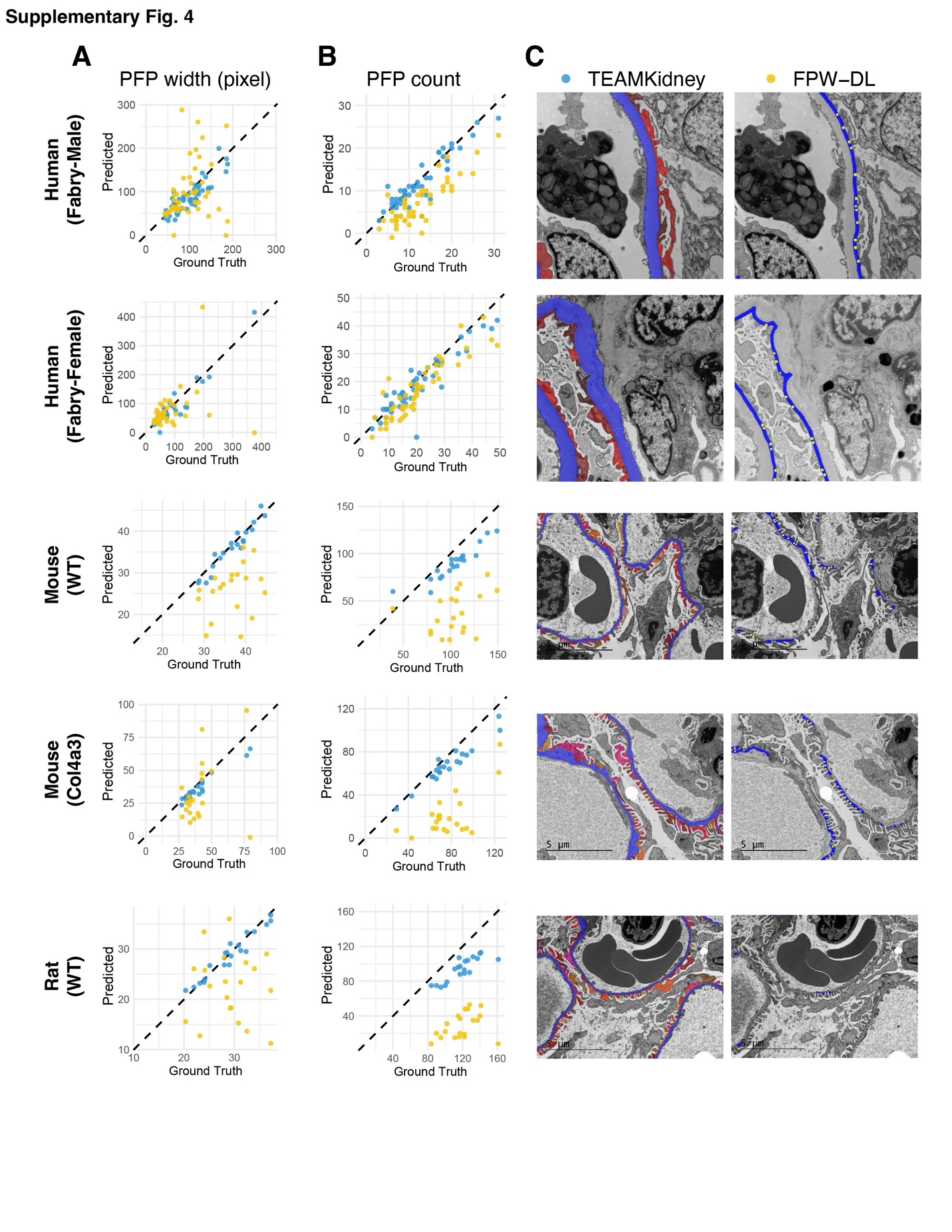

**Supplementary Fig. 4.** **Comparison of FP quantification between TEAMKidney and FPW-DL methods in additional models.** Scatter plots with diagonal reference lines show the agreement between model predictions and ground truth measurements for (A) average PFP width and (B) PFP counts. (C) Representative TEM images with segmentation masks generated by TEAMKidney (left) and FPW-DL (right). From top to bottom: male and female Fabry disease patients, WT mouse model, Col4a3 mouse model and WT rat model.

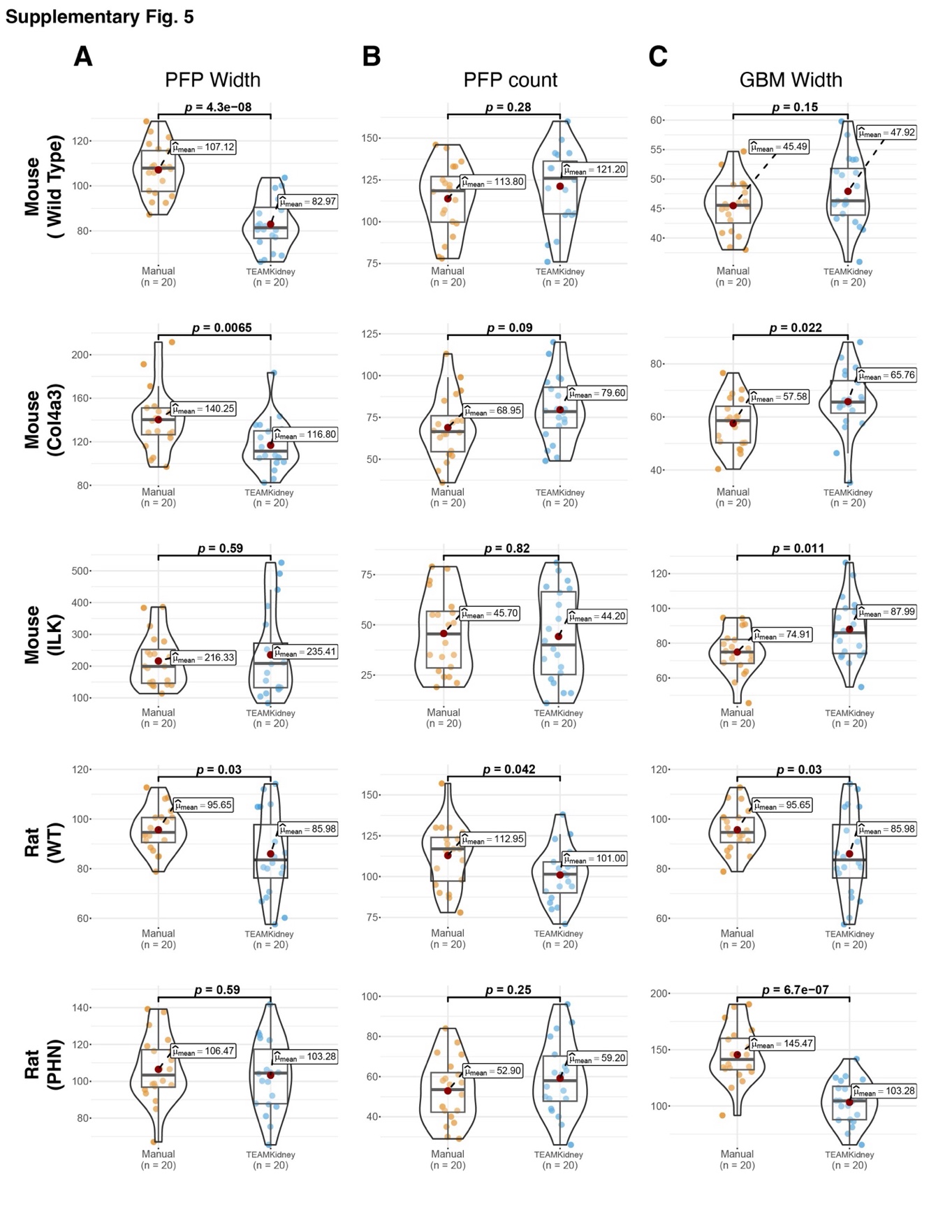
**Supplementary Fig. 5.** **Evaluation of concordance between expert manual measurements and TEAMKidney predictions on animal models.** (A) GBM mean width. (B) FP mean width, (C) FP count. From top to bottom: WT mouse, Col4a3 mouse, ILK mouse, WT rat, and PHN rat model.

**Supplementary Table 1.** **Performance comparison between the proposed Glom2Mask model and the benchmark Mask2Former for panoptic segmentation.**

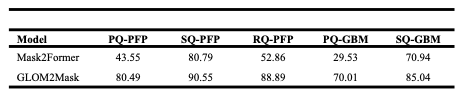
